# Pawpaws prevent predictability: A locally-dominant tree alters understory beta-diversity and community assembly

**DOI:** 10.1101/2024.03.04.583351

**Authors:** Anna C. Wassel, Jonathan A. Myers

**Affiliations:** Department of Biology, Washington University in St. Louis, St. Louis, Missouri, 63130, USA

**Keywords:** *Asimina triloba*, beta-diversity, community assembly, community size, competition, dominant species, ecological drift, forest herbs, null model, species interactions, stochasticity, temperate forest

## Abstract

While dominant species are known to be important in ecosystem functioning and community assembly, biodiversity responses to the presence of dominant species can be highly variable. Dominant species can increase the importance of deterministic community assembly by competitively excluding species in a consistent way across local communities, resulting in low site-to-site variation in community composition (beta-diversity) and non-random community structure. In contrast, dominant species could increase the importance of stochastic community assembly by reducing the total number of individuals in local communities (community size), resulting in high beta-diversity and more random community structure. We tested these hypotheses in a large, temperate oak-hickory forest plot containing a locally-dominant tree species, pawpaw (*Asimina triloba*; Annonaceae), an understory tree species that occurs in dense, clonal patches in forests throughout the east-central United States. We determined how the presence of pawpaw influences local species diversity, community size, and beta-diversity by measuring the abundance of all vascular plant species in 1x1-m plots both inside and outside pawpaw patches. To test whether the presence of pawpaw influences local assembly processes, we compared observed patterns of beta-diversity inside and outside patches to a null model of random assembly. We found lower local species diversity, lower community size, and higher observed beta-diversity inside pawpaw patches than outside pawpaw patches. Moreover, standardized effect sizes of beta-diversity from the null model were lower inside pawpaw patches than outside pawpaw patches, indicating more random community composition inside pawpaw patches. Together these results suggest that pawpaw increases the importance of stochastic relative to deterministic community assembly at local scales, likely by decreasing overall numbers of individuals, and increasing random local extinctions inside patches. Our findings provide insights into the ecological processes by which locally-dominant tree species shape the assembly and diversity of understory plant communities at different spatial scales.

## Introduction

Highly abundant species within communities can have strong effects on biodiversity and ecosystem functioning (Grime 1998, Gaston 2011, Avolio et al. 2019). Species that have high abundance relative to other species in a community *and* proportionate effects on environmental conditions, community diversity, and/or ecosystem functioning are considered “dominant species” (Avolio et al. 2019). Dominant species can determine nutrient cycling and primary productivity (Grime 1998, Ellison 2019), increase resistance or resilience of ecosystems to environmental change (Avolio et al. 2019), add physical structure to a habitat (Dayton 1972, Altieri and Witman 2014), and modify the abiotic environment in ways that create more harsh conditions or conversely ameliorate abiotic stress for other species (Hughes 2010, Lustenhouwer et al. 2012, Gavilán and Callaway 2017). Although the loss of dominant species can have cascading effects on communities and ecosystems, their effects on patterns of species diversity can be highly variable (Myers and Harms 2009, Hughes 2010, Gavilán and Callaway 2017, Avolio et al. 2019, Ellison et al. 2019, Elsberry and Bracken 2021). This variation potentially reflects multiple ecological processes through which dominant species affect community assembly, but the relative roles of these processes remain understudied.

Dominant species can affect community assembly through deterministic or stochastic processes. Deterministic processes include abiotic filtering and biotic interactions such as competition and facilitation that reflect niche differences among species in a community (Vellend 2010, Chase and Myers 2011, Leibold and Chase 2017). Dominant species can increase the importance of interspecific competition when they limit space or resources for other species (e.g. Lustenhouwer et al. 2012), resulting in competitive exclusion (Konno 2002, Segre et al. 2014, Ellison et al. 2015). Alternatively, dominant species can facilitate the survival of certain species by lowering abiotic stress (e.g. Gavilán and Callaway 2017). Dominant species can also increase the importance of stochastic community assembly by decreasing the total number of individuals in a local community (local community size) (Powell et al. 2013). As local community size decreases, more species in the community may become rare, thereby increasing demographic stochasticity and random changes in species’ relative abundances (ecological drift; MacArthur and Wilson 1967, Hubbell 2001, Orrock and Watling 2010). In addition, amelioration of stressful conditions by dominant species can lead to more random assembly of nondominant species (Arnillas and Cadotte 2019). The effects of dominant species on deterministic and stochastic processes are expected to increase when dominant species are also of large stature, i.e., when size asymmetries among competing species or guilds are large (Keddy and Shipley 1989, Myers and Harms 2009).

Despite widespread interest in the role of dominant species in communities and ecosystems (Ellison et al. 2005, Gilbert et al. 2009, Avolio et al. 2019), their relative effects on deterministic and stochastic community assembly remain unresolved. First, previous studies have largely focused on how dominant species influence species diversity at local spatial scales (e.g., alpha diversity), but similar patterns of local species diversity could reflect different assembly processes. For example, low species richness can result from either competitive exclusion by dominant species (Konno 2002, Segre et al. 2014, Ellison et al. 2015) or random local extinctions in small communities with few individuals (Powell et al. 2013). Most dominant-species removal experiments in plant communities have focused on changes in local species richness or diversity, finding a mix of positive (Konno 2002, Segre et al. 2014, Ellison et al. 2015, Avolio et al. 2019), negative (Hughes 2010, Altieri and Witman 2014, Gavilán and Callaway 2017), or no clear response (Myers and Harms 2009, Gilbert et al. 2009) to the removal of dominant plant species. Second, relatively few studies have examined how dominant species influence site-to-site variation in community composition (beta-diversity). Patterns of beta-diversity can help elucidate the relative importance of deterministic and stochastic processes (Anderson et al. 2011, Chase and Myers 2011). For example, deterministic exclusion of inferior competitors by dominant species should cause local communities to converge in composition (i.e., low beta-diversity), whereas random local extinctions in small communities should cause local communities to diverge in composition (i.e., high beta-diversity). Finally, observed changes in beta-diversity can be compared to a null model of random community assembly to further assess the relative roles of deterministic and stochastic processes (Chase 2007, Catano et al. 2017). Therefore, patterns of diversity at different scales can provide key insights into the ecological roles of dominant species in community assembly and ecosystem functioning.

In this study, we examined the effect of a locally-dominant tree species, pawpaw (*Asimina triloba*; Annonaceae), on the diversity and assembly of understory plant communities in a temperate forest-dynamics plot. Our focal species, pawpaw, is a widely-distributed understory tree species that occurs in dense, clonal patches in forests throughout the east-central United States. Pawpaw has been shown to be a dominant species in temperate forests with high local abundance (Appendix S1: Fig. S1) and strong effects on the diversity of other tree species (Baumer and Runkle 2010). While the assembly of forest tree communities is fairly well studied (e.g. Condit et al. 2000, Condit et al. 2002, Ellison et al. 2019), the assembly of forest herb communities has received less attention, despite the disproportionate contribution of herbaceous plant species to temperate forest diversity (Gilliam 2007, Spicer et al. 2020). We therefore examined the effect of pawpaw on both the total understory community (woody and herbaceous species combined) and herbaceous species only.

We tested two non-mutually exclusive hypotheses. First, we tested the hypothesis that pawpaw increases the relative role of deterministic assembly through interspecific competition (hereafter the deterministic assembly hypothesis). Second, we tested the hypothesis that pawpaw increases the relative role of stochastic assembly by decreasing local community size (hereafter the stochastic assembly hypothesis). The deterministic assembly hypothesis predicts that the presence of dominant species 1) decreases local species diversity due to competitive exclusion, 2) decreases beta-diversity among local communities by selecting for a limited subset of species that can co-occur with dominant species, and 3) results in lower beta-diversity than expected by random assembly from the species pool. In contrast, the stochastic assembly hypothesis predicts that the presence of dominant species 1) decreases local species diversity, but 2) increases beta-diversity among local communities, due to random local extinctions, and 3) results in beta-diversity that is more similar to patterns expected under random assembly from the species pool. We tested these predictions by comparing observed patterns of local species diversity, local community size, and beta-diversity among paired groups of understory plant communities located inside and outside of pawpaw patches. We then compared observed patterns of beta-diversity to a null model that simulated random assembly of local communities from the species pool.

## Methods

### Study site and focal species

We conducted this study at Washington University in St. Louis’ environmental field station, Tyson Research Center, located 25 miles from St. Louis, Missouri. The 800-ha site is located on the edge of the Ozark highlands, dominated by late-successional, deciduous oak-hickory forest, and contains a topographically heterogeneous landscape characterized by silty loam and silty clay soils that develop from shale and cherty limestone (Zimmerman and Wagner 1979). Our study was conducted within the Tyson Research Center Forest Dynamics Plot, a large (20.16 ha; 480 x 420 m), stem-mapped forest plot that is part of the Forest Global Earth Observatory (ForestGEO) network (Anderson-Teixeira et al. 2015). The 20-ha plot includes more than 1,600 stems of pawpaw at least 1 cm in diameter at breast height (DBH), most of which occur in 18 patches ranging in area from 5-1028 m^2^.

Our focal dominant species for this study is the pawpaw tree, *Asimina triloba* (Annonaceae) (hereafter pawpaw). Pawpaw is distributed widely throughout the east-central United States and parts of southern Canada (Sullivan 1993), making it the northernmost member of the otherwise tropical family Annonaceae. It primarily occurs in moist valleys and mesic hillsides (Immel and Anderson 2001). Pawpaw can reproduce both sexually and asexually, forming dense, discrete clonal patches (Hosaka et al. 2005). While not the most abundant species in temperate forests at larger spatial scales due to its patchy distribution, at our study site it is frequently the most abundant species at the 10×10 m scale when it is present (Appendix S1: Fig. S1), making it a *locally* dominant species. The local dominance and discrete patch structure of this species make it an ideal study system for investigating how the presence or absence of a dominant species affects community assembly processes.

### Sampling design

We selected five blocks to contain a pawpaw patch and an adjacent area without pawpaws, referred to as “inside” and “outside” patches, respectively (Fig. 1, Fig. 2). The inside (pawpaw) patches selected ranged from 58-435 m^2^ in size (mean = 189 m^2^). The paired outside patches were selected to have abiotic (soil and topographic) conditions similar to those inside the pawpaw patch and were 10 to 20 m from the edge of the pawpaw patch (Fig. 2a, 2c). We determined the similarity of soil and topographic conditions between the inside and outside patches through a Principal Component Analysis (PCA) on 17 soil and topographic variables (Appendix S1: Fig. S2). The values were estimated for each 10 _J 10-m subplot in the 20-ha ForestGEO plot based on measurements taken in 2013 (detailed in Spasojevic et al. 2014, LaManna et al. 2016). The outside patches were chosen to have a similar PC1 score as the pawpaw patches.

**Fig. 1.**
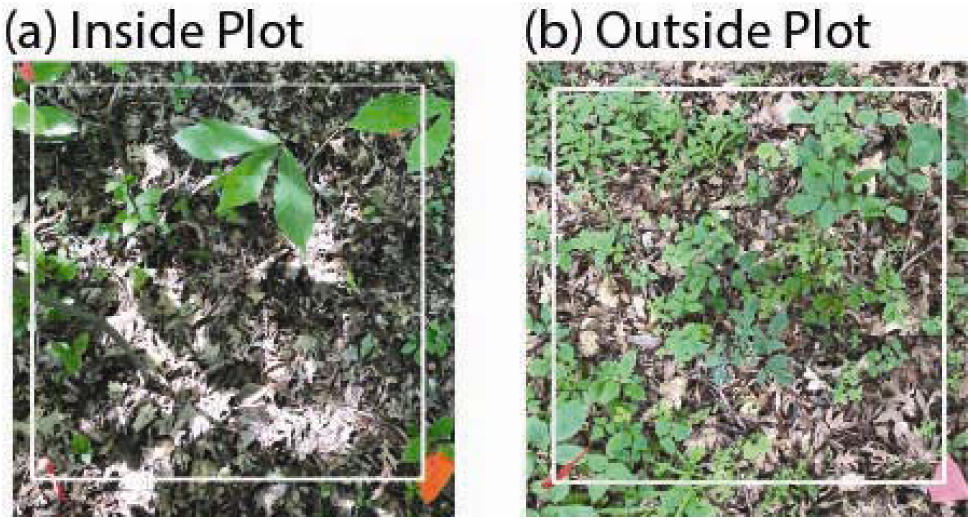
Examples of understory plant communities (a) inside a pawpaw (*Asimina triloba*) patch (“inside plot”) and (b) outside a pawpaw patch at least 10 m away from the patch edge (“outside plot”). White squares show 1×1-m plots. Photos by Anna C. Wassel.

**Fig. 2.**
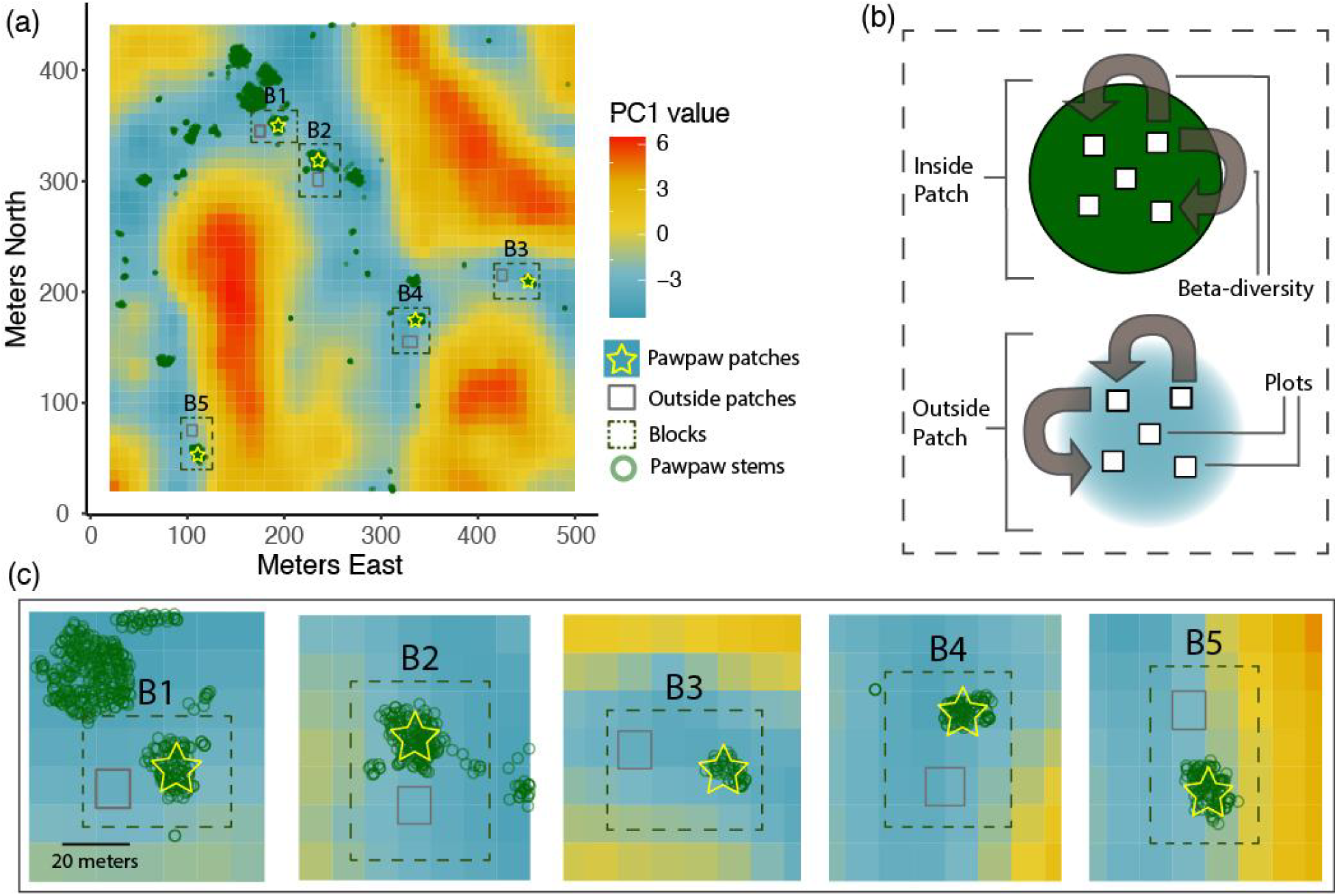
Sampling design within the Tyson Research Center ForestGEO Plot, Missouri. (a) Abiotic conditions (soil resources & topography) represented by the first axis of a principal component analysis (PCA) including 17 environmental variables at the 10×10-m scale (Appendix S1: Fig. S2), mapped locations of all pawpaw stems ≧1 cm in diameter at breast height (DBH), and selected sampling blocks. Blue values represent areas of lower elevation, higher soil-nutrient availability, and higher soil pH, whereas red values represent higher elevation, lower nutrient availability, and more acidic soils. (b) For each block, the pawpaw patch edge was defined and five 1×1-m plots were placed inside or outside the patch. Gray arrows represent how beta-diversity was calculated within each of the two patch types (inside and outside plots). (c) Each block is shown zoomed in to illustrate the environmental conditions and relative location for both the inside patches (yellow stars over pawpaw stems) and outside patches represented by gray rectangles.

For each patch type in each of the five blocks, five 1×1-m plots were sampled for plant community composition (n=25 inside plots, n=25 outside plots, n=50 plots total; Fig. 2b). Each plot was at least two m within the edge of the pawpaw patch but did not contain any woody stems over 1 cm DBH. Woody stems over 1 cm DBH were excluded to keep standard the amount of available ground area available for understory plants. We recorded the identity and estimated the abundance of all understory vascular plant species, i.e. herbaceous plants including ferns, and woody plants and vines. We estimated abundance (number of stems per species) as the number of 10×10-cm cells that contained rooted stems of the given species. In cases when individuals could not be identified to species in the field, they were identified to genus or assigned a morphospecies and photos were uploaded to iNaturalist for future assistance with identification; 8% of stems were considered morphospecies for analyses. We estimated local community size by summing the abundances of all species in each plot. Sampling was conducted during the peak growing season from July to September of 2021. Due to the different life stages and biology of young woody seedlings compared to the adult herbaceous plants, we conducted two separate analyses for: 1) herbaceous and woody plant species combined (hereafter total understory community); and 2) herbaceous species only.

### Analyses

We analyzed local species diversity, community size, and beta-diversity using linear mixed-effects models in R (package ‘nlme’; Pinheiro et al. 2023). All models included patch type (inside or outside) as a fixed effect and block as a random effect. When necessary, we log-transformed response variables to meet the assumptions of homogeneous variances between patch types and normality of model residuals. When transformation did not improve homogeneity of variances, we used a heterogeneous variance model (‘varIdent’ function). We describe the analyses for each response variable below.

To test our first prediction, we calculated local species diversity using the inverse Simpson’s index (Simpson 1949, Oksanen et al. 2022); a scale-independent diversity measure of the effective number of species that is insensitive to differences in numbers of individuals (Chase et al. 2018). For the model testing local diversity of the total understory community, we log-transformed the inverse-Simpson’s values to meet the assumption of homogeneous variances. For the model testing local diversity of herbaceous species only, we used a heterogeneous variance model and excluded the two plots with no species.

To test our second prediction, we calculated observed beta-diversity as the compositional dissimilarity among plots using the Bray-Curtis index. We analyzed beta-diversity based on distance-to-centroid values (Anderson 2006, Kraft et al. 2011) using the ‘betadisper’ function in the R vegan package (Oksanen et al. 2022), where each value represents the distance (compositional dissimilarity) from an individual plot to the centroid of the group of all 25 plots within each patch type (Fig. 2b). When analyzing beta-diversity of herbaceous species only, we excluded two inside plots from block 5 that contained no herbaceous plants.

To test our third prediction, we used a null model to simulate the compositional dissimilarity expected by random community assembly (Kraft et al. 2011, Myers et al. 2013, LaManna et al. 2021). First, we defined the species pool as all species recorded during the study across all inside and outside plots combined. We estimated the total abundance of each species (number of stems) in the species pool by summing its frequencies (number of 10×10-cm cells in which a rooted stem was recorded) across all plots. Second, in each of 2000 iterations of the null model, we simulated community assembly in each plot by randomly sampling stems from the species pool, while keeping constant the empirically observed total number of stems in each plot (local community size) and total abundance of each species in the species pool. Third, we calculated the mean simulated beta-diversity for each plot by averaging the Bray-Curtis distance-to-centroid values from the 2000 null-model iterations. Fourth, we calculated the standardized effect size as the difference between the observed beta-diversity (distance-to-centroid) and mean simulated values for each plot, divided by the standard deviation of simulated values for each plot. A standardized effect size of zero indicates that observed beta-diversity does not differ from random sampling of the species pool, a positive value indicates higher beta-diversity than expected by chance, and a negative value indicates lower beta-diversity than expected by chance. We tested median standardized effect sizes of each patch type against the null expectation of zero with one-sample two-sided Wilcoxon tests. All analyses were conducted in R (R Core Team 2022).

## Results

Overall, we observed a total of 79 plant species and morphospecies (hereafter species) in this study, including 52 herbaceous plant species and 27 woody plant species (Appendix S1: Table S1, S2, S3). Only 6 species were unique to inside patches while there were 29 species unique to outside patches. Of the 52 herbaceous plant species, 24 occurred inside pawpaw patches and 47 occurred in outside patches. Of the 27 woody plant species, 12 occurred inside pawpaw patches and 26 occurred in outside patches. Of the species that occurred in both patch types (inside and outside), most had lower abundance inside pawpaw patches (Appendix S1: Fig. S3). Among taxa identified to the species level (non-morphospecies, Appendix S1: Table S1, S2), herbaceous species were more abundant than woody species outside of pawpaw patches (68.4% of the total estimated number of stems), but less abundant than woody species inside pawpaw patches (39.9% of the total estimated number of stems).

### Local species diversity and community size

Local species diversity and community size were significantly lower inside than outside pawpaw patches (Fig. 3, Appendix S1: Table S4). For herbaceous species only, median local diversity was 49% lower inside than outside pawpaw patches (Fig. 3a). Median community size (total estimated number of rooted stems of all species in a plot) for herbaceous species was 76% lower inside than outside pawpaw patches (Fig. 3b). Similar patterns were observed for the total understory community (herbaceous and woody species combined). For the total understory community, median local diversity was 29% lower inside than outside pawpaw patches (Fig. 3a), and median community size was 67% lower inside than outside pawpaw patches (Fig. 3b).

**Fig. 3.**
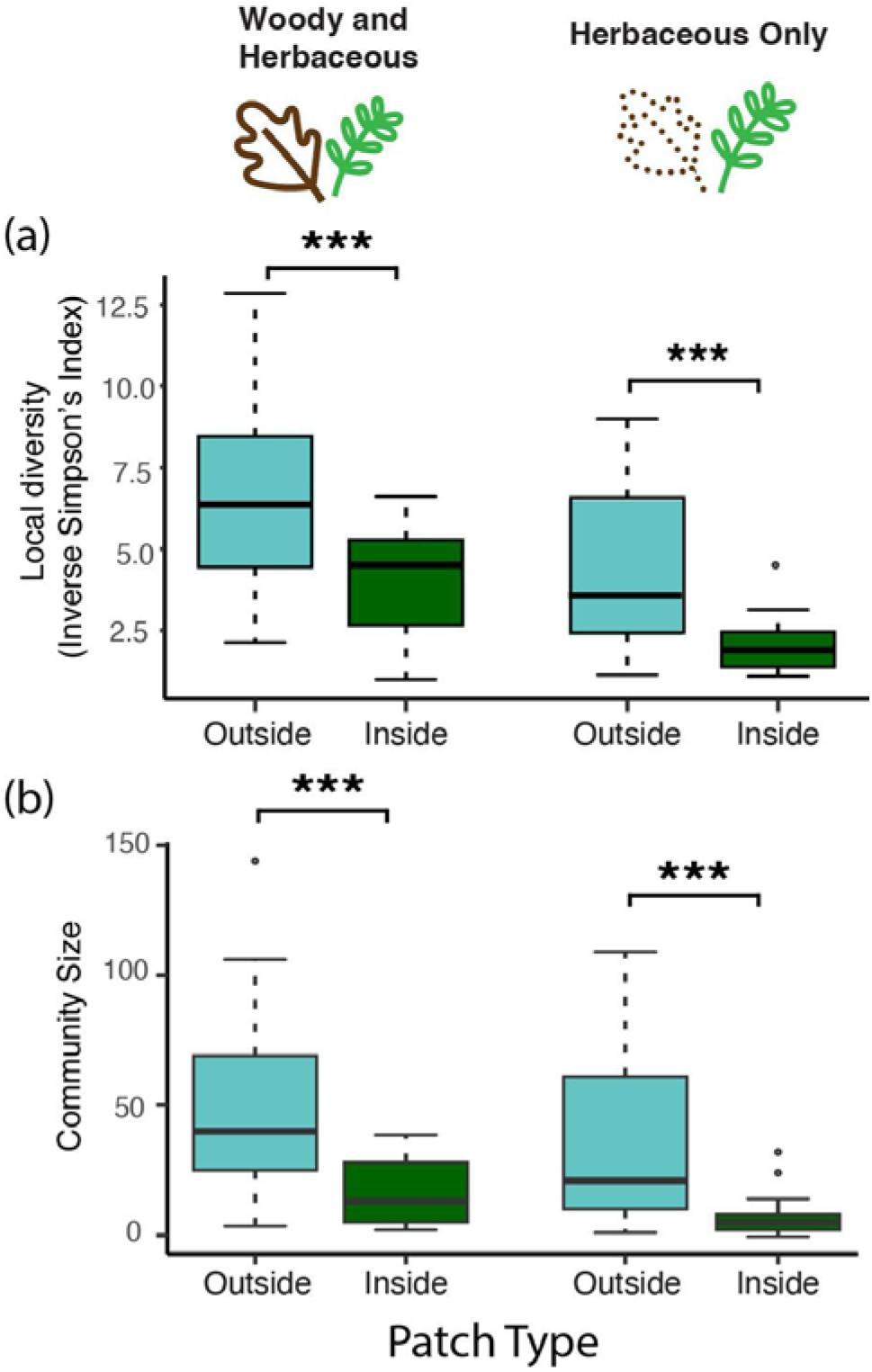
Local plant species diversity and community size are lower inside than outside of pawpaw patches. (a) Local species diversity (inverse Simpson’s index) outside (N = 25) and inside plots (N = 25) for the total understory community (herbaceous and woody plant species), and herbaceous species only. (b) Local community size for the total understory community, and herbaceous species only. Community size was estimated as the total number of rooted stems of all species in each plot. Boxes represent the median and 25th/75th percentile, whiskers extend to the largest value up to 1.5 times the interquartile range, and dots represent outlier data points. (*** = *P* < 0.001)

### Observed, simulated, and standardized effect sizes of beta-diversity

Observed, simulated, and standardized effect sizes of beta-diversity differed significantly inside and outside pawpaw patches for herbaceous species only (Fig. 4a-c; Appendix S1: Table S4). Observed and simulated beta-diversity was higher inside than outside pawpaw patches (Fig. 4a-b). In contrast, standardized effect sizes of beta-diversity were significantly lower inside than outside pawpaw patches (Fig. 4c). Median standardized effect sizes inside and outside of pawpaw patches were both positive and differed significantly from zero, though the difference was less significant inside pawpaw patches (Fig. 4c; Appendix S1: Table S5; *P* = 0.044 inside patches; *P* = 0.001 outside patches). Similar patterns were observed for the total understory community, with the exception of observed beta-diversity, which showed no significant difference between patch types. Median standardized effect sizes differed more between patch types, due to larger standardized effect sizes outside of pawpaw patches for the total understory community (Fig. 4f) compared to herbaceous species only (Fig. 4c). For the total understory community (Fig. 4f), median standardized effect sizes inside and outside pawpaw patches both differed significantly from zero (Table S4; *P* = 0.013 inside patches; *P* < 0.001 outside patches).

**Fig. 4.**
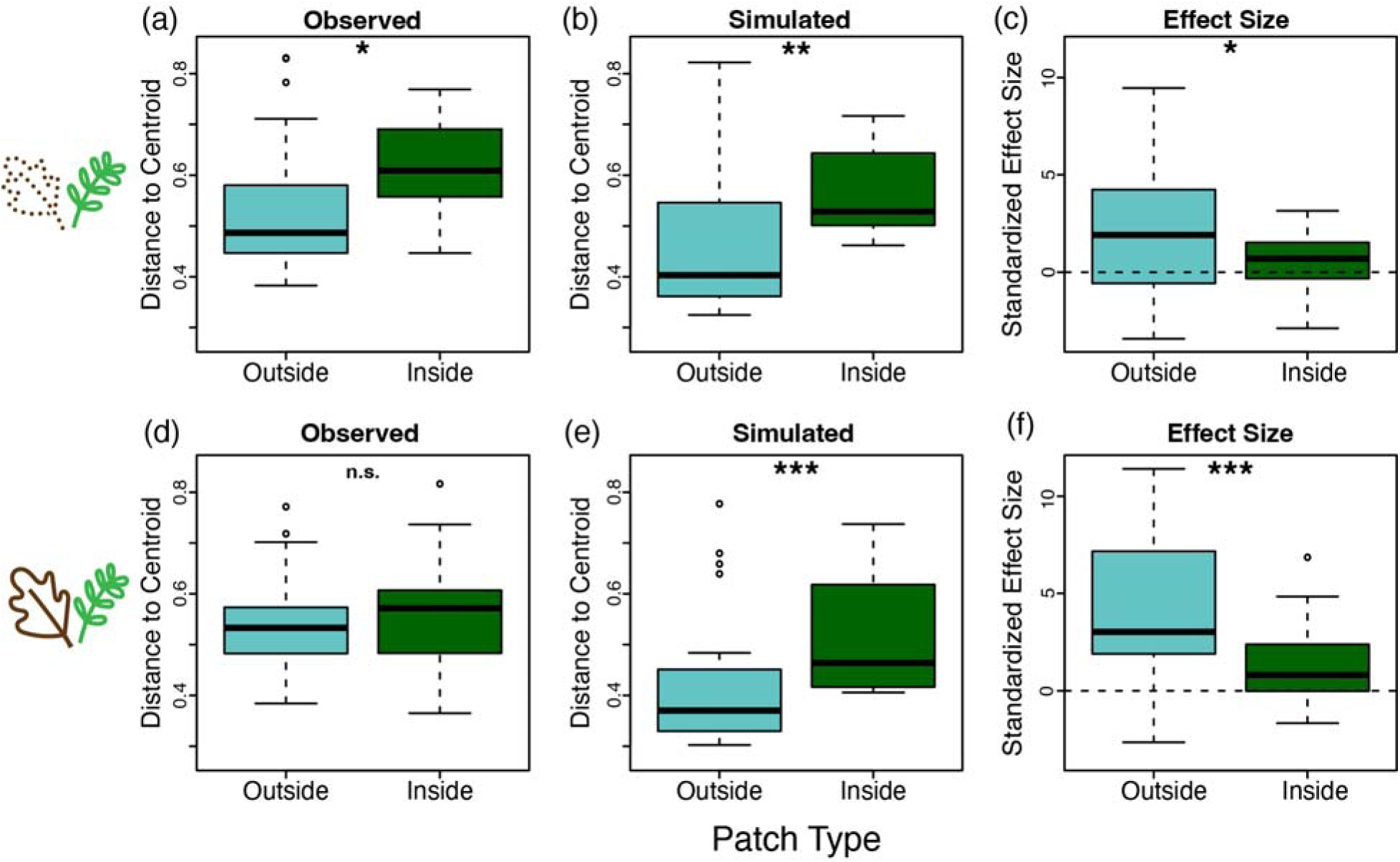
Variation in species composition (beta-diversity) differs inside and outside of pawpaw patches. (a) Observed beta-diversity of herbaceous plant species in plots outside (N = 25) and inside (N = 23) pawpaw patches. (b) Simulated beta-diversity expected from a null model of random assembly. (c) Standardized effect size of beta-diversity. Panels (d), (e), and (f) show the same results including herbaceous and woody species. The dashed line at zero represents the null expectation. Boxes represent the median and 25th/75th percentile, whiskers extend to the largest value up to 1.5 times the interquartile range, and dots represent outlier data points. (n.s. = not significant, * = *P* < 0.05, ** = *P* < 0.01, *** = *P* < 0.001)

## Discussion

Overall, our results support the stochastic assembly hypothesis. The lower local species diversity, lower community size, and more random variation in species composition found within pawpaw patches all support predictions of the stochastic assembly hypothesis. Beta-diversity was higher than expected by chance both inside and outside of pawpaw patches, but patterns of beta-diversity inside pawpaw patches more closely resembled the null expectation of random community assembly. These findings suggest that both deterministic and stochastic assembly processes are important in shaping the understory plant community, but that local communities in different patch types are not assembled the same way, with stochastic processes being relatively more important than deterministic processes in patches dominated by pawpaw.

Observed beta-diversity of herbaceous species was higher inside than outside of pawpaw patches, supporting the second prediction of the stochastic assembly hypothesis (Fig. 4a). Higher beta-diversity among plots inside pawpaw patches is in line with the findings of other studies that observed the presence of dominant woody species increases beta-diversity (Ellison et al. 2015, Ellison et al. 2019,) or decreases local relative to regional diversity (Powell et al. 2013). However, previous studies have often focused on how dominant tree species affect other tree species, without explicit consideration of their effects on herbaceous species. In our study, the difference in observed beta-diversity of herbaceous species inside and outside of pawpaw patches (Fig. 4a) became weaker and non-significant when considering the total understory community of herbaceous and woody species together (Fig. 4d). This indicates that abundances of woody species are consistent enough across the patch types to reduce overall differences in community composition. This could be due to several factors. First, woody species are generally less dispersal-limited than herbaceous species (Turnbull et al. 2000). In our study, for example, the most-common species of woody seedlings present inside pawpaw patches was northern spicebush (*Lindera benzoin*) (Appendix S1: Table S1), a bird-dispersed shrub with high adult abundance at our site. With increased dispersal, we expect decreased beta-diversity (Leibold and Chase 2017, Germain et al. 2017, Ron et al. 2018, Thompson et al. 2020). Second, the woody seedlings are at a life stage that experiences higher mortality and is generally less diverse than the adult tree community (Green et al. 2014, Ramachandran et al. 2023). Seedling communities have been shown to have lower beta-diversity than adult tree communities (Ramachandran et al. 2023), presumably due to these ontogenetic differences in the importance of different assembly mechanisms (Cavender-Bares and Bazzaz 2000, Comita et al. 2007, Green et al. 2014, Spasojevic et al. 2014). Meanwhile, the herbaceous community largely contains diverse adult assemblages that survived past the seedling stage. Lastly, most of the plant species diversity in temperate forests is comprised of herbaceous species, many of which are rare in the understory (Gilliam 2007, Spicer et al. 2020), such that including more common woody species will inherently shift the composition towards being more similar. These results illustrate the need to further investigate how herbaceous communities assemble in forests, as differences in functional diversity and life-stages between herbaceous and woody species can elucidate different assembly processes in the understory (Spicer et al. 2022).

Patterns of beta-diversity inside pawpaw patches more closely resembled the null expectation of random assembly (Fig. 4c, f), supporting the third prediction of the stochastic assembly hypothesis.. The smaller effect sizes inside pawpaw patches could reflect several ecological processes. First, theory (Hubbell 2001, Orrock and Watling 2010, Vellend 2016) and empirical studies (e.g. Gilbert and Levine 2017, Siqueira et al. 2020, Fodelianakis et al. 2021) show that decreases in community size causes random changes in species relative abundances (ecological drift), thereby increasing compositional variation among local communities. In our study, community size was 49–76% lower inside than outside pawpaw patches (Fig. 3b), and of the species present in both patch types, most had much lower abundance inside pawpaw patches (Appendix S1: Fig. S3), likely making local populations within pawpaw patches more prone to demographic stochasticity. Second, simulation models exploring the interplay between selection and ecological drift show that high beta-diversity can emerge when niche-based processes exacerbate the effects of neutral processes (Latombe et al. 2015). For example, Gilbert and Levine (2017) demonstrated that the presence of a dominant competitor can deterministically lower other species’ relative abundances to a point where stochasticity has an even greater effect, causing very high species turnover among their plots with the smallest overall community sizes. Third, larger null-model deviations outside of pawpaw patches can reflect more deterministic processes such as fine-scale environmental heterogeneity, local plant-soil and plant-plant interactions, and species-specific differences in dispersal ability (Condit et al. 2002, Bauer et al. 2017, Germain et al. 2017, Thompson et al. 2020). These deterministic processes likely contribute to the non-random patterns of beta-diversity observed inside and outside pawpaw patches, as well as the relatively stronger non-random patterns observed outside pawpaw patches.

We also found that local species diversity was consistently lower inside than outside pawpaw patches (Fig. 3a). Previous studies have found that dominant plant species can decrease local diversity (e.g. Myers and Harms 2009, McCain et al. 2010, Ellison et al. 2015, Hejda et al. 2019, Hernández et al. 2022, Eckberg et al. 2023), but the underlying ecological processes remain unresolved. Although our study cannot discern the degree to which low species diversity inside patches reflects dispersal limitation, non-random competitive exclusion, or ecological drift, lower community size may increase the role of ecological drift inside pawpaw patches. The effects of community size and dispersal limitation may be further exacerbated in larger pawpaw patches, where dispersal from source populations located outside patches may be less likely to balance local extinctions of dispersal-limited herbs inside pawpaw patches. Additionally, our findings are in contrast to studies that found that some dominant species facilitated species diversity by mitigating harsh conditions, often at the edge of subordinate species’ range (Dayton 1972, Pellissier et al. 2010, Gavilán and Callaway 2017, Elsberry and Bracken 2021).

Several abiotic and biotic factors may explain the lower community size, lower local species diversity, and more random patterns of beta-diversity within pawpaw patches. First, above and below ground abiotic conditions may be altered by pawpaw trees. Pawpaws have been shown to be strong competitors for light which could decrease the abundances of otherwise shade-tolerant understory plants (Cole and Weltzin 2005). In addition, high pawpaw stem densities and clonal growth may increase belowground competition for soil nutrients and water (Baumer and Runkle 2010). Second, pawpaw may be allelopathic (McEwan et al. 2010, Pavliuchenko 2018). In our study ecosystem, sites invaded by the allelopathic shrub, bush honeysuckle (*Lonicera mackii*), have low diversity of native plant species (Powell et al. 2013), making this a particularly intriguing hypothesis. However, the current evidence for allelopathy in pawpaws is weak (McEwan et al. 2010, Pavliuchenko 2018) to negative (Cole and Weltzin 2005). Third, pawpaw’s interaction with a dominant herbivore, white-tailed deer (*Odocoileus virginianus*), may explain patterns of diversity. Pawpaw is unpalatable to deer, leading deer to selectively browse other species (Slater and Anderson 2014, Shelton et al. 2014, Jenkins et al. 2015). If deer are selectively browsing the herbaceous layer in pawpaw communities to avoid the unpalatable pawpaw leaves, this could decrease community size, decrease local species diversity, and increase beta-diversity within pawpaw patches. Alternatively, if deer avoid pawpaw patches altogether due to their inedibility, this could potentially decrease seed dispersal by deer via endozoochory and epizoochory of new propagules into pawpaw patches (e.g. Myers et al. 2004, Blyth et al. 2013, Guiden 2017). Finally, a combination of suboptimal niche conditions and medium to high dispersal rates may make local communities within pawpaw patches subject to source-sink dynamics, with pawpaw patches harboring “sink” populations (Pulliam 1988).

Our study highlights several avenues for future research on the mechanisms by which pawpaw shapes forest community assembly. Future studies can use seed-addition experiments to test the degree to which low species diversity (Myers and Harms 2009) and high beta-diversity (Germain et al. 2017) of herbaceous species are caused by dispersal limitation within pawpaw patches. Similarly, investigating the degree to which micro-habitat variation in light and soil conditions inside and outside pawpaw patches determine community composition can help further differentiate between deterministic and stochastic assembly processes. Future studies can also explore how pawpaw patch characteristics such as patch size, age, and demography affect the strength of these processes and biodiversity patterns. Long-term studies of woody plant recruitment, growth, and survival inside and outside of pawpaw patches can elucidate how pawpaws may affect forest regeneration (Baumer and Runkle 2010, Hochwender et al. 2016) or invasive species spread (Cole and Weltzin 2005). Further understanding the biology and ecology of this and other locally-dominant tree species will provide key insights into how species interactions drive the assembly, diversity, and dynamics of understory plant communities at varying spatial scales.

## Supporting information

AppendixS1

## Acknowledgements

We thank Erin O’Connell, Nathan Aaron, Aspen Workman, James Lucas, Noah Dell, and Brad Delfeld for species identification support, Sean W. McHugh for coding assistance and coffee, Christopher Catano for sharing his species list and mapping code, members of the Myers Lab, Landis Lab and Sebastian Tello for discussions and comments on the manuscript, the Tyson Research Center staff, and the more than 140 research technicians, undergraduate students, and high school students that have contributed to the Tyson Research Center Forest Global Earth Observatory (ForestGEO) Plot Project. This project was supported by a Webster Groves Nature Study Society (WGNSS) Bo Koster Scholarship to ACW, George Hayward Plant Biology Graduate Fellowship to ACW, Maxwell/Hanrahan Foundation Field Work Grant from the Missouri Botanical Garden to ACW, National Science Foundation grants DEB 1557094 and DEB 2240431 to JAM, the International Center for Advanced Renewable Energy and Sustainability (I-CARES) at Washington University in St. Louis, ForestGEO, Washington University in St. Louis’ Provost’s Office, and Tyson Research Center.

