## AppendixS1 for "Pawpaws prevent predictability: A locally-dominant tree alters understory beta-diversity and community assembly"

**Appendix S1**

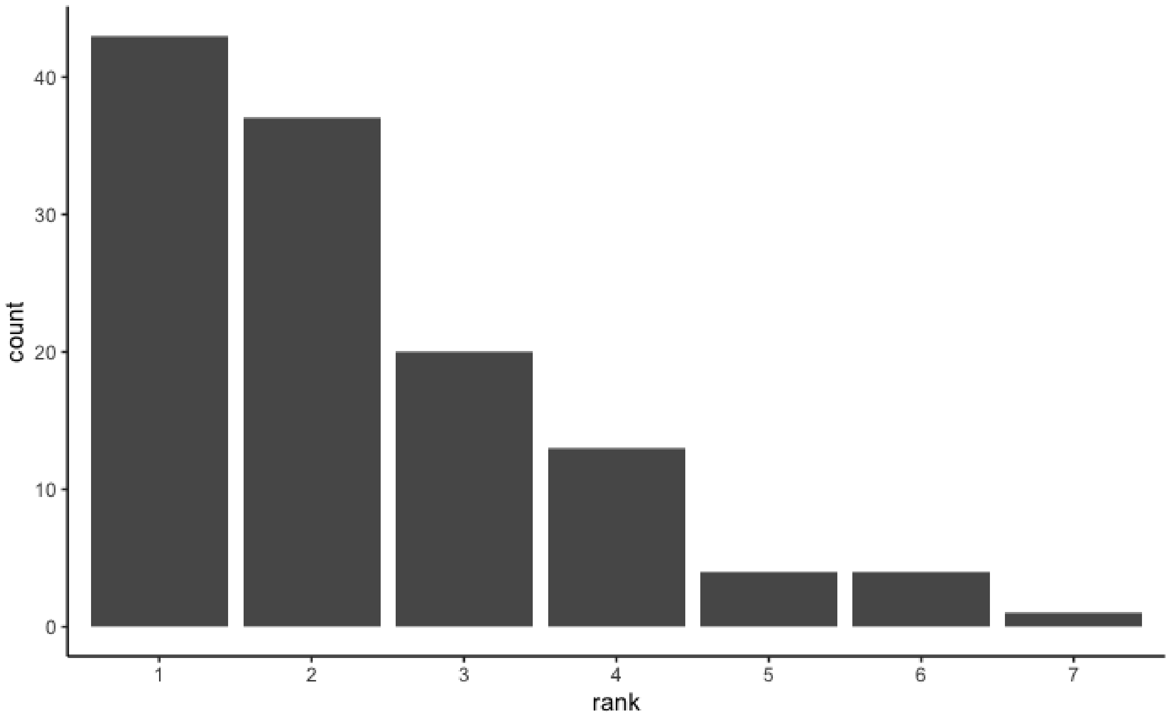


Fig. S1. Rank abundance of pawpaw (*Asimina triloba*) in every 10x10 meter subquadrat in which at least one stem ≥1 cm in diameter at breast height (DBH) is present at the Tyson Research Center Forest Dynamics Plot. The plurality of occurrences of pawpaw are in areas where it is the most abundant species.

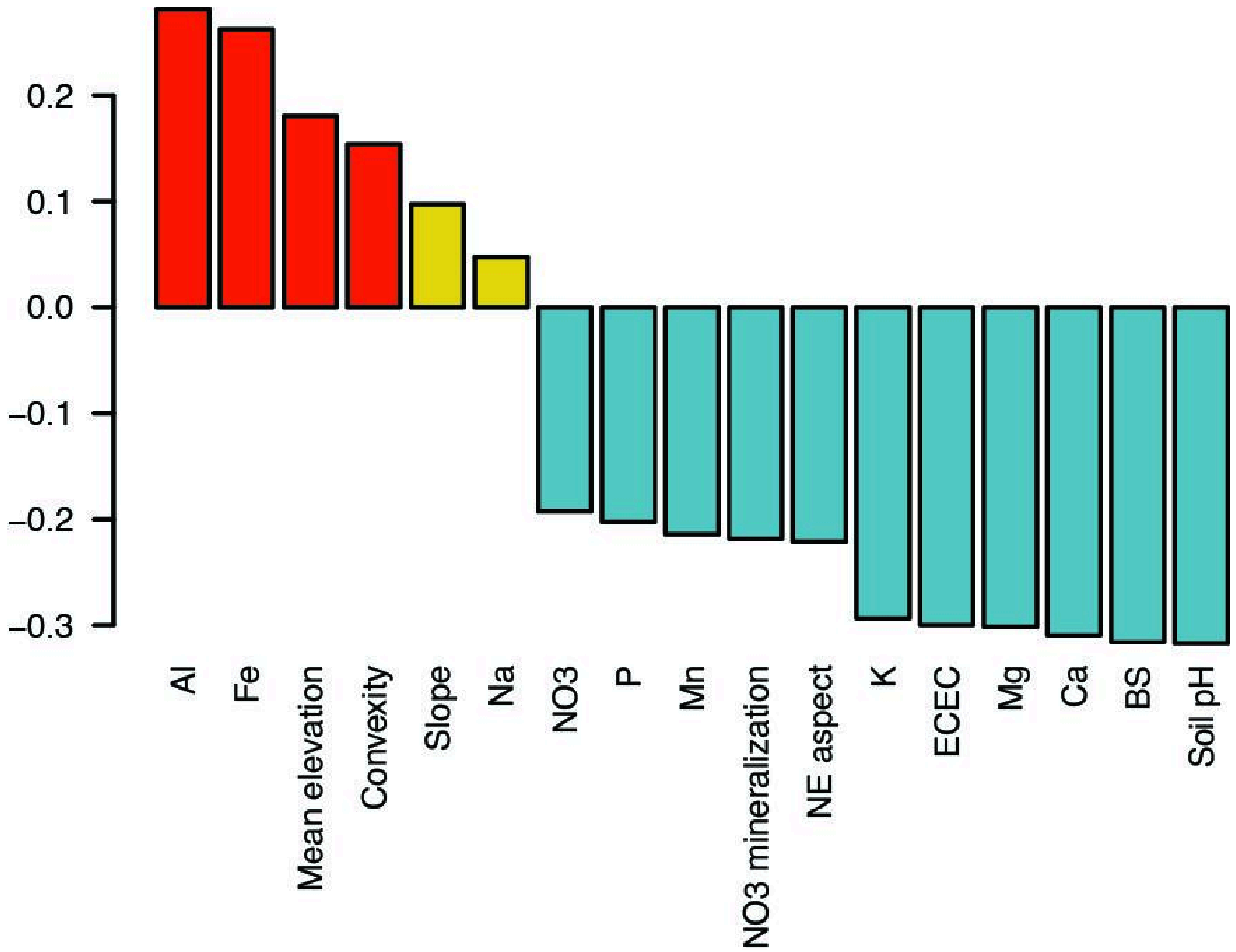


Fig. S2. Loadings from the first axis of a principal component analysis (PCA) of 17 soil and topographic variables in the Tyson Research Center Forest Dynamics Plot. Al, Fe, Na, P, Mn, K, Mg, and Ca refer to soil nutrients measured in mg/kg. $NO_3$ min is the nitrate mineralization rate over ~10 days (mg/kg). ECEC is Effective Cation Exchange Capacity ($cmol_c$/kg) calculated as: Al+Ca+Fe+K+Mg+Mn+Na. BS is base saturation (%) calculated as: (Ca+K+Mg+Na) / (Al+Ca+Fe+K+Mg+Mn+Na). Mean elevation (m), terrain convexity (m), slope (degrees), and northeastern (NE) aspect (radians) refer to topographic variables. Colors correspond to the colors of PC1 values as shown in Fig. 2.

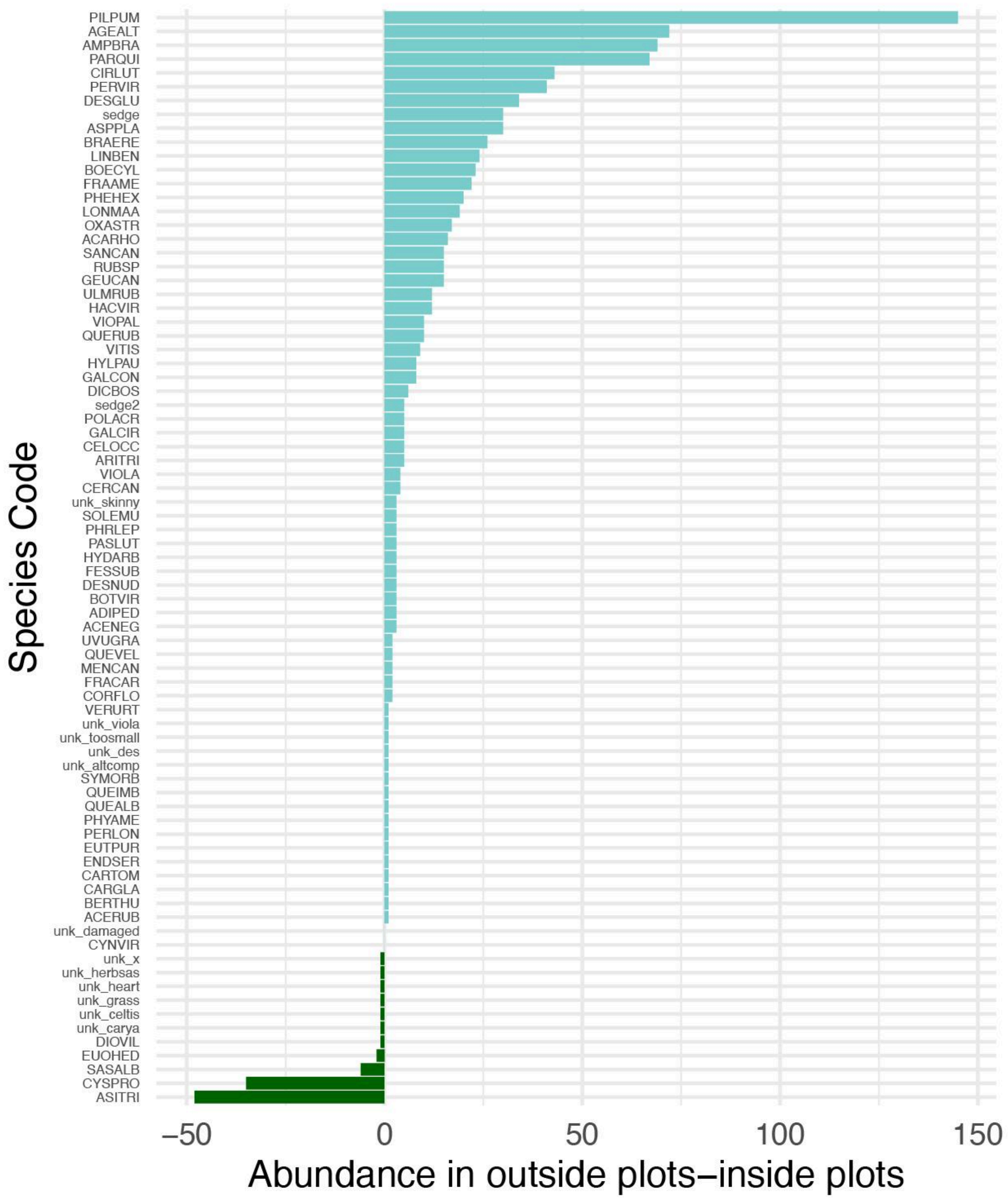


Fig. S3. The difference in species' abundance across all blocks between outside and inside plots. Positive values in light blue indicate the species was more abundant outside pawpaw patches overall. See Tables S1, S2, and S3 for full species names.

Table S1 Woody Species List

| Species Code | Genus | Species | Family | Total abun-dance | Inside abun-dance | Outside abun-dance |
|---|---|---|---|---|---|---|
| LINBEN | *Lindera* | *benzoin* | Lauraceae | 174 | 75 | 99 |
| PARQUI | *Parthenocissus* | *quinquefolia* | Vitaceae | 155 | 44 | 111 |
| LONMAA | *Lonicera* | *maackii* | Caprifoliaceae | 65 | 23 | 42 |
| ASITRI | *Asimina* | *triloba* | Annonaceae | 58 | 53 | 5 |
| FRAAME | *Fraxinus* | *americana* | Oleaceae | 34 | 6 | 28 |
| ULMRUB | *Ulmus* | *rubra* | Ulmaceae | 24 | 6 | 18 |
| SASALB | *Sassafras* | *albidum* | Lauraceae | 22 | 14 | 8 |
| QUERUB | *Quercus* | *rubra* | Fagaceae | 10 | 0 | 10 |
| ACENEG | *Acer* | *negundo* | Sapindaceae | 9 | 3 | 6 |
| CERCAN | *Cercis* | *canadensis* | Fabaceae | 8 | 2 | 6 |
| CELOCC | *Celtis* | *occidentalis* | Cannabaceae | 7 | 1 | 6 |
| HYDARB | *Hydrangea* | *arborescens* | Hydrangeaceae | 3 | 0 | 3 |
| QUEALB | *Quercus* | *alba* | Fagaceae | 3 | 1 | 2 |
| CORFLO | *Cornus* | *florida* | Cornaceae | 2 | 0 | 2 |
| EUOHED | *Euonymus* | *hederaceus* | Celastraceae | 2 | 2 | 0 |
| MENCAN | *Menispermum* | *canadense* | Menispermaceae | 2 | 0 | 2 |
| QUEVEL | *Quercus* | *velutina* | Fagaceae | 2 | 0 | 2 |
| ACERUB | *Acer* | *rubra* | Sapindaceae | 1 | 0 | 1 |
| BERTHU | *Berberis* | *thunbergii* | Berberidaceae | 1 | 0 | 1 |
| CARGLA | *Carya* | *glabra* | Juglandaceae | 1 | 0 | 1 |
| CARTOM | *Carya* | *tomentosa* | Juglandaceae | 1 | 0 | 1 |
| QUEIMB | *Quercus* | *imbricaria* | Fagaceae | 1 | 0 | 1 |
| SYMORB | *Symphoricarpos* | *orbiculatus* | Caprifoliaceae | 1 | 0 | 1 |

Table S2 Herbaceous Species List

| Species Code | Genus | Species | Family | Total abun-dance | Inside abun-dance | Outside abun-dance |
|---|---|---|---|---|---|---|
| PILPUM | *Pilea* | *pumila* | Urticaceae | 203 | 29 | 174 |
| CIRLUT | *Circaea* | *lutetiana* | Onagraceae | 107 | 32 | 75 |
| AGEALT | *Ageratina* | *altissima* | Asteraceae | 82 | 5 | 77 |
| AMPBRA | *Amphicarpaea* | *bracteata* | Fabaceae | 71 | 1 | 70 |
| CYSPRO | *Cystopteris* | *protrusa* | Dryopteridaceae | 71 | 53 | 18 |
| PERVIR | *Persicaria* | *virginiana* | Polygonaceae | 47 | 3 | 44 |
| DESGLU | *Desmodium* | *glutinosum* | Fabaceae | 38 | 2 | 36 |
| ASPPLA | *Asplenium* | *platyneuron* | Aspleniaceae | 30 | 0 | 30 |
| HACVIR | *Hackelia* | *virginiana* | Boraginaceae | 30 | 9 | 21 |
| BRAERE | *Brachyelytrum* | *erectum* | Poaceae | 26 | 0 | 26 |
| BOECYL | *Boehmeria* | *cylindrica* | Urticaceae | 23 | 0 | 23 |
| SANCAN | *Sanicula* | *canadensis* | Apiaceae | 21 | 3 | 18 |
| PHEHEX | *Phegopteris* | *hexagonoptera* | Thelypteridaceae | 20 | 0 | 20 |
| OXASTR | *Oxalis* | *stricta* | Oxalidaceae | 19 | 1 | 18 |
| ACARHO | *Acalypha* | *rhomboidea* | Euphorbiaceae | 18 | 1 | 17 |
| PHRLEP | *Phryma* | *leptostachya* | Phrymaceae | 17 | 7 | 10 |
| GEUCAN | *Geum* | *canadense* | Rosaceae | 15 | 0 | 15 |
| VIOPAL | *Viola* | *palmata* | Violaceae | 10 | 0 | 10 |
| GALCON | *Galium* | *concinnum* | Rubiaceae | 8 | 0 | 8 |
| HYLPAU | *Hylodesmum* | *pauciflorum* | Fabaceae | 8 | 0 | 8 |
| DICBOS | *Dichanthelium* | *boscii* | Poaceae | 6 | 0 | 6 |
| FRACAR | *Frangula* | *carolinana* | Rhamnaceae | 6 | 2 | 4 |
| ARITRI | *Arisaema* | *triphyllum* | Araceae | 5 | 0 | 5 |
| BOTVIR | *Botrypus* | *virginanus* | Ophioglossaceae | 5 | 1 | 4 |
| DIOVIL | *Dioscorea* | *villosa* | Dioscoreaceae | 5 | 3 | 2 |
| GALCIR | *Galium* | *circaezans* | Rubiaceae | 5 | 0 | 5 |
| POLACR | *Polystichum* | *acrostichoides* | Dryopteridaceae | 5 | 0 | 5 |
| ADIPED | *Adiantum* | *pedatum* | Pteridaceae | 3 | 0 | 3 |
| DESNUD | *Desmodium* | *nudliflorum* | Fabaceae | 3 | 0 | 3 |
| FESSUB | *Festuca* | *subverticillata* | Poaceae | 3 | 0 | 3 |
| PASLUT | *Passiflora* | *lutea* | Passifloreaece | 3 | 0 | 3 |
| SOLEMU | *Solanum* | *emulans* | Solanaceae | 3 | 0 | 3 |
| CYNVIR | *Cynoglossum* | *virginianum* | Boraginaceae | 2 | 1 | 1 |
| UVUGRA | *Uvularia* | *grandiflora* | Liliaceae | 2 | 0 | 2 |
| ENDSER | *Endodeca* | *serpentaria* | Aristolochia | 1 | 0 | 1 |
| EUTPUR | *Eutrochium* | *purpureum* | Asteraceae | 1 | 0 | 1 |

| | | | | | | |
|---|---|---|---|---|---|---|
| PERLON | *Persicaria* | *longiseta* | Polygonaceae | 1 | 0 | 1 |
| PHYAME | *Phytolacca* | *americana* | Phytolaccaceae | 1 | 0 | 1 |
| VERURT | *Verbena* | *urticifolia* | Verbenaceae | 1 | 0 | 1 |

Table S3 List of plants in study that were identified to genus and morphospecies.

| Species Code | Genus | Species | Family | Total abun-dance | Inside abun-dance | Outside abun-dance |
|---|---|---|---|---|---|---|
| sedge | unknown | unknown | unknown | 46 | 8 | 38 |
| VITIS | *Vitis* | unknown | Vitaceae | 35 | 13 | 22 |
| RUBSP | *Rubus* | unknown | Rosaceae | 15 | 0 | 15 |
| unk_altcomp | unknown | unknown | unknown | 9 | 4 | 5 |
| sedge2 | unknown | unknown | unknown | 5 | 0 | 5 |
| VIOLA | *Viola* | unknown | Violaceae | 4 | 0 | 4 |
| unk_damaged | unknown | unknown | unknown | 4 | 2 | 2 |
| unk_carya | *Carya* | unknown | Juglandaceae | 3 | 2 | 1 |
| unk_skinny | unknown | unknown | unknown | 3 | 0 | 3 |
| unk_celtis | *Celtis* | unknown | Cannabaceae | 1 | 1 | 0 |
| unk_des | unknown | unknown | unknown | 1 | 0 | 1 |
| unk_grass | unknown | unknown | Poaceae | 1 | 1 | 0 |
| unk_heart | unknown | unknown | unknown | 1 | 1 | 0 |
| unk_herbsas | unknown | unknown | unknown | 1 | 1 | 0 |
| unk_toosmall | unknown | unknown | unknown | 1 | 0 | 1 |
| unk_viola | unknown | unknown | Violaceae | 1 | 0 | 1 |
| unk_x | unknown | unknown | unknown | 1 | 1 | 0 |

Table S4: Results from linear mixed-effects models testing for effect of patch type (inside or outside a pawpaw patch) on local species diversity (Inverse Simpson's Index), local community size (total estimated number of rooted stems of all species in a plot), and beta-diversity (Bray Curtis distance-to-centroid) of the total understory community (herbaceous and woody species) and herbaceous species only. Standardized effect size = (observed beta-diversity - mean simulated beta-diversity) / standard deviation of simulated beta-diversity. ***$P \leq 0.001$; **$P \leq 0.01$, *$P < 0.05$

| Response variable | Test statistic (F value) | Degrees of freedom (denDF) | P value | Significance level |
|---|---|---|---|---|
| Local species diversity | | | | |
| Herbaceous species only | 4.47 | 42 | 0.0001 | *** |
| Total understory community | 4.61 | 44 | 0.0001 | *** |
| Local community size | | | | |
| Herbaceous species only | 26.91 | 44 | <.0001 | *** |
| Total understory community | 33.79734 | 44 | <.0001 | *** |
| Observed beta-diversity | | | | |
| Herbaceous species only | 6.0703 | 42 | 0.0179 | * |
| Total understory community | 1.14241 | 44 | 0.291 | n.s. |
| Simulated beta-diversity | | | | |
| Herbaceous species only | 7.76375 | 42 | 0.008 | ** |
| Total understory community | 5.08259 | 44 | 0.0003 | *** |
| Standardized effect size | | | | |
| Herbaceous species only | 5.387344 | 42 | 0.0252 | * |
| Total understory community | 14.29977 | 44 | 0.0005 | *** |

Table S5. Results from one-sample Wilcoxon tests for standardized effect sizes of beta-diversity inside and outside of pawpaw patches compared to $\mu = 0$. ***$P \leq 0.001$; **$P \leq 0.01$, *$P < 0.05$

| Response variable | Plant Community | Patch type | Test statistic (V) | P value | Significance level |
|---|---|---|---|---|---|
| Standardized effect size | Herbaceous only | Inside | 204 | 0.0448 | * |
| Standardized effect size | Herbaceous only | Outside | 275 | 0.0016 | ** |
| Standardized effect size | Total understory | Inside | 253 | 0.0135 | * |
| Standardized effect size | Total understory | Outside | 314 | <0.0001 | *** |
